# Multi-compartment impact of micropollutants and particularly antibiotics on bacterial communities using environmental DNA at river basin-level

**DOI:** 10.1101/2024.03.28.587215

**Authors:** Pedro A. Inostroza, Gerdhard L. Jessen, Feilong Li, Xiaowei Zhang, Werner Brack, Thomas Backhaus

## Abstract

Microbial communities, in particular bacterial assemblies, play pivotal roles in sustaining biogeochemical processes within ecosystems. They are also responsible for the degradation of toxic chemicals, while the development of resistance against antimicrobial drugs jeopardises human health. Bacterial communities respond to environmental conditions with diverse structural and functional changes depending on their compartment (water, biofilm or sediment), type of environmental stress, and type of pollution to which they are exposed. In this study, we combined amplicon sequencing of bacterial 16S rRNA genes from water, biofilm, and sediment samples collected in the anthropogenically impacted River Aconcagua basin (Central Chile, South America), in order to evaluate whether micropollutants alter bacterial community structure and functioning based on the type and degree of chemical pollution. Furthermore, we evaluated the potential of bacterial communities from differently polluted sites to degrade contaminants. Our results show a lower diversity at sites impacted by agriculture and urban areas, featuring high loads of micropollution with pesticides, pharmaceuticals and personal care products as well as industrial chemicals. Nutrients, antibiotic stress, and micropollutant loads explain most of the variability in the sediment and biofilm bacterial community, showing a significant increase of bacterial groups known for their capabilities to degrade various organic pollutants, such as *Nistrospira* and also selecting for taxa known for antibiotic resistance such as *Exiguobacterium* and *Planomicrobium*. Moreover, potential ecological functions linked to the biodegradation of toxic chemicals at the basing level revealed significant reductions in ecosystem-related services in sites affected by agriculture and wastewater treatment plant (WWTP) discharges across all investigated environmental compartments. Finally, we suggest transitioning from simple concentration-based assessments of environmental pollution to more meaningful toxic pressure values in order to comprehensively evaluate the role of micropollutants at the ecological (biodiversity) level.

**Graphical abstract:** 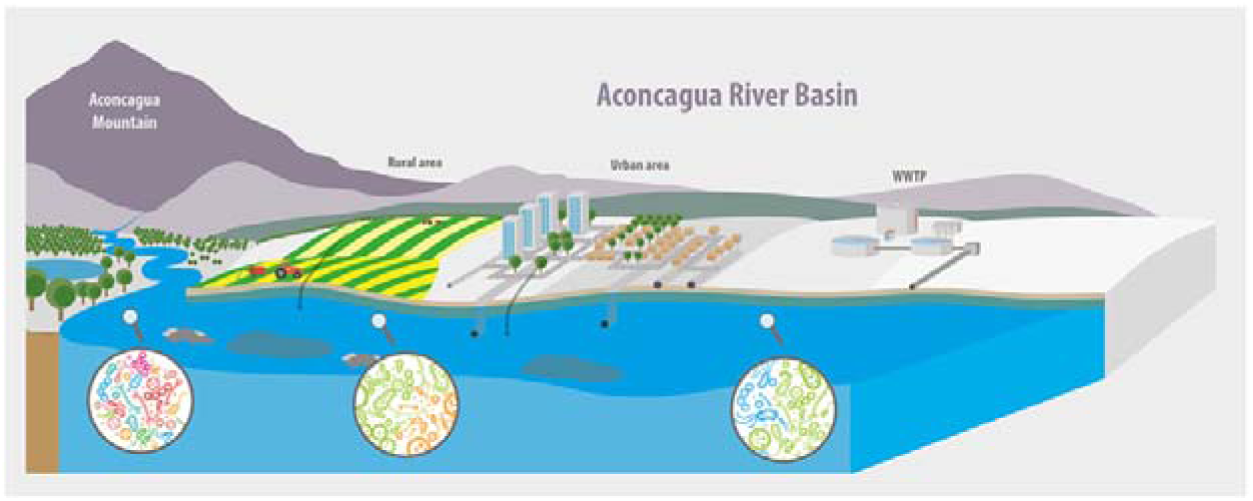

**Highlights:**

- Micropollutant mixtures altered bacterial community structure and functioning
- Antibiotic stress correlated significantly with changes in community structure
- Reduction of ecological functions related to the degradation of contaminants
- Water, biofilm, and sediments relevant for microbial ecotoxicology

## 1. Introduction

Anthropogenic chemical pollution with pesticides, pharmaceuticals, personal care products, industrial chemicals and other synthetic organic chemicals, stands as one of the major drivers affecting aquatic biodiversity (Sigmund et al., 2023). Such chemicals enter the aquatic environment as a consequence of their inefficient removal in domestic and industrial WWTPs (Hug et al., 2014; Kaewlaoyoong et al., 2018), as well as via runoff from agricultural (Reiber et al., 2021), and urban areas (Inostroza et al., 2023a). Although many micropollutants are not persistent, their continuous release into the aquatic environment, renders many of them “pseudo-persistent” (Boxall et al., 2004; Kolpin et al., 2002). Synthetic organic chemicals are typically present at low environmental concentrations in the aquatic environment (pg/L to ng/L) and are therefore often called micropollutants (Schwarzenbach et al., 2006). Micropollutants often have deleterious effect at different trophic levels (Beketov et al., 2013; Burdon et al., 2020; Kidd et al., 2007; Parrish et al., 2019) and levels of biological organisation (Inostroza et al., 2016; König et al., 2017; Weichert et al., 2020) due to their high biological activity (Stamm et al., 2016).

Microbial communities, especially bacterial communities, are key players that drive many of the biogeochemical processes responsible of sustaining ecosystems (Falkowski et al., 2008). They provide crucial ecosystem services, including the degradation of toxic chemicals (Aguilar-Romero et al., 2020). Furthermore, bacterial communities develop or adopt resistant mechanisms (Davies and Davies, 1996; Rizzo et al., 2013) due to long-term exposure to antimicrobial chemicals, which threatens public health worldwide.

The responses of bacterial community to micropollutants are complex, due to their site-specific composition and, therefore, resilience, as well as their functional redundancy. For instance, Burdon et al. (2020) found alterations in the bacterial community structure downstream of WWTPs, marked by significant compositional changes and reduced biodiversity. Similar responses were observed in bacterial communities in sediments downstream of WWTPs (Drury et al., 2013; Freixa et al., 2020). Studies that consider broader spatial distributions have also reported a reduced diversity of bacterial communities that inhabit areas characterised by urban and agricultural land use, compared to reference areas (Ibekwe et al., 2016; Li et al., 2023). However, it is important to note that other studies that compared sites up- and downstream of WWTPs did not find significant biodiversity differences (Mansfeldt et al., 2020) and changes in bacterial community composition in areas of agricultural and industrial land use were attributed to chemical pollution in water and sediment (Valverde et al., 2021). Indeed, significant associations have been described between micropollutant concentrations and alterations in bacterial communities in water samples, but not in sediments in mixed land use areas (Gao et al., 2019).

Despite their ecological significance, microbial communities are often ignored in the assessment of ecological quality (Brandt et al., 2015). Even strict environmental legislative frameworks, such as the European Union Water Framework Directive (EU WFD), do not include microbial communities as one of their Biological Quality Elements. Furthermore, even if pollution impacts on bacterial communities are assessed, the analysis often relies on correlations with measured environmental concentrations, ignoring the vastly differing toxicity profiles of micropollutants. A more meaningful description of pollution impacts is, however, possible by normalising concentrations to the toxicity of the chemicals encountered, making use of the well-established “toxic unit” concept (Sprague, 1970). In the end, the combined use of chemical-analytical measurements with culture-independent methods, such as DNA- and/or RNA-sequencing-based approaches, might be the most promising approach to shed light on alterations of the microbial biodiversity caused by chemical pollution.

In this study, we introduce a novel approach to comprehensively evaluate structural (i.e., alpha- and beta-diversity), compositional, and functional changes of bacterial communities in response to chemical pollution. The overall objectives were to: i) characterise bacterial community structure in different environmental compartments, including surface waters, biofilm, and superficial sediments using amplicon sequencing of bacterial 16S rRNA genes; ii) explore associations between toxicity- normalised micropollutant concentrations and bacterial communities to infer the impact of micropollutants on structure and potential function of bacterial communities and; iii) evaluate potential alterations in bacterial ecological functions due to micropollutants in each environmental compartment and land use context.

We conducted our study in the River Aconcagua basin in Central Chile (South America), as a model for an anthropogenically impacted river basin (Fierro et al., 2021; Fuentes et al., 2015; Inostroza et al., 2023a, 2024). The microbial communities inhabiting this ecosystem are subjected to runoff from intensive agricultural activities, micropollutant discharges from several WWTPs (Inostroza et al., 2023a) located in urban and rural areas, and intensive industrial activities associated with metallic mining (mainly copper).

## 2. Material and methods

### 2.1. Case study and chemical analyses – River Aconcagua Basin

The River Aconcagua Basin, situated in Central Chile, has a drainage area of 7,338 km^2^. The climate in the basin is Mediterranean, with dry and warm Austral summers (October to April) and cool, wet winters (May to September), which are often characterised by irregular and intense precipitation events (Amigo and Ramírez, 1998). The basin has a population of ca. half a million people and supports more than 10% of Chile’s national agricultural and 4% of its copper production, respectively (COCHILCO, 2020). Briefly, agriculture accounts for almost 8% of the total river basin area, with over 90% of the cropland, comprised by avocado and grape farms, concentrated in the lower sub-basins (Webb et al., 2021). Additionally, there are ten medium sized WWTPs located throughout the basin to treat wastewater for almost half a million residents. Five of these WWTPs discharge directly into the main course of the River Aconcagua, while the remaining five discharge into tributaries. Detailed information on the wastewater treatment technologies used, location, and population served can be found in Table S1.

In October 2018 (warm/dry season), environmental DNA (eDNA) was collected from nine sampling sites based on land use (Figure S1). Thus, three sites, designated as “reference sites,” were characterised by low urban/chemical pressure and located in middle mountain areas (RS1 and RS2) and within a national park (RS3). Three other sites were situated in small tributary streams running throughout agricultural areas (T1 and T3) and mixed land uses (T2). Lastly, three sites were located along the main course of the River Aconcagua (R1, R2, and R3). This land use classification is supported by pollution profiles and environmental risk assessment works. These studies highlight low micropollution loads in reference sites, middle micropollution loads in tributaries, and the highest concentrations in the main river course (Inostroza et al., 2023a). Sites at ecological risk due to micropollutants are found in tributaries and the main river course (Inostroza et al., 2024). A detailed description of the sampling sites is provided in Table S2.

At each site, *in situ* measurements of physicochemical parameters, including temperature, pH, conductivity, dissolved oxygen, and light intensity were conducted using a multi-parameter probe (Hannah model HI 9829, United Kingdom) or HOBO data loggers (Onset, USA). To assess main nutrients, 15 mL surface water was filtered in triplicate through a 0.45 µm GF/F glass fibre filter and stored at −20□°C until laboratory analysis. Nitrate (NO^3-^), nitrite (NO^2-^), phosphate (PO_4_^-3^), and silicic acid (SiOH_4_^-4^) were measured using standard colourimetry methods, with a segmented flow Seal AutoAnalyser (Seal Analytical AA3).

### 2.2. Sampling eDNA for bacterial communities

To characterise the bacterial communities, cells were collected from surface water, biofilm, and surface sediment in all nine sites in the River Aconcagua Basin (Figure S1). For the water bacterial community, cells were retrieved from four independent 250 mL water samples at each site and filtered through sterile 0.22 µm pore size hydrophilic nylon membranes (Nalgene, Thermo Scientific). Biofilm cells were collected from at least five cobblestones per site (epilithic biofilm) by brushing the biofilms off using sterilised toothbrushes. Then, samples were merged and transferred to 50 mL centrifuge tubes and fixed with ethanol (final concentration 70%) until DNA extraction. Finally, superficial sediment (top 0-5 cm) was collected using sterile 50 mL single-use tip-off syringes and transferred to sterile plastic bags. All samples were kept in dark at −20 °C until DNA extraction.

### 2.3. DNA extraction protocols, library preparation, sequencing, and bioinformatics pipelines

DNA from water, biofilm, and sediment was extracted using the DNeasy PowerWater Kit, DNeasy PowerBiofilm Kit, and DNA PowerSoil Pro Kit, respectively (Qiagen, Germany), following a modified version of the manufacturer’s instructions in the case of filters (Chiang and Inostroza, 2022). The V3 region of 16S rRNA genes were amplified using the Bacteria-specific primer pair 341F (ACCTACGGGRSGCWGCAG) and 518R (GGTDTTACCGCGGCKGCTG; Shanghai Generay Biotech Co., Ltd.) described by Klindworth et al. (2012).

In accordance with Li et al. (2018), triplicate polymerase chain reaction (PCR) reactions were executed for each sample to diminish potential PCR bias. The PCR reactions were conducted in a 50 μL reaction mixture that contained 31 μL of ddH_2_O, 10 μL of 5× Phusion Green HF Buffer, 1 μL of 10 mM dNTPs, 2.5 μL of each primer (10 μM), 2.5 μL of DNA template, and 0.5 μL of Phusion Green Hot Start II High-Fidelity DNA Polymerase (Thermo Fisher). The amplification protocol consisted of an initial denaturation step at 98 °C for 30 s, followed by 30 cycles at 98 °C for 5 s, 62 °C for 30 s, and 72 °C for 15 s, with a final extension at 72 °C for 7 min. The PCR products were then assessed by visualising on a 2% agarose gel to ensure that the expected size of PCRs was yielded. Following visualisation, the PCR products were purified using the E-Z 96 Cycle Pure Kit (Omega) and quantified with Qubit dsDNA HS Assay Kits (Invitrogen). The purified PCR products were then pooled equally for subsequent sequencing. To prepare sequencing templates, sequencing adaptors were linked to the purified DNA fragments with the Ion Xpress Plus Fragment Library Kit (Thermo Fisher) in accordance with the manufacturer’s protocol. Unique 12-nt nucleotide tags were attached to the 5’-ends of the forward or reverse primers to pool all samples for optimised sequencing. Finally, all samples were diluted to a final concentration of 100 pM, and sequencing templates were generated using Ion OneTouch 2 and sequenced in the Ion Proton sequencer (Life Technologies).

The split_libraries.py script in the QIIME toolkit was employed to eliminate raw sequence data of low quality, defined as having a mean quality score less than 20, sequences with ambiguous “N”, homopolymers, and sequence lengths of less than 100 bp (Caporaso et al., 2010). After cleaning, the reads were sorted and distinguished using unique sample tags, and subsequently subjected to OTU clustering at a cut-off value of 97% nucleotide similarity, following the UPARSE pipeline as described by Edgar (2013). For taxonomic assignment of each OTU, the Greengenes database (DeSantis et al., 2006) was used in conjunction with the align_seqs.py script.

### 2.4. Microbial community analysis

The operational taxonomic unit (OTU) data was analysed using the R-package phyloseq (McMurdie and Holmes, 2013). Unassigned annotations together with OTUs identified as “chloroplast” and “mitochondria” were removed. Raw reads were scrutinised for inferior performance during amplification, pooling, or sequencing steps that resulted in contamination or sample loss. Alpha diversity based on Observed Richness, Chao1, Evenness, and Shannon diversity metrics were calculated for each sample before and after rarefaction based on Knight et al. (2018) suggestions. No rarefied data is presented since no differences were found between the two datasets. To compare alpha diversities, Kruskal-Wallis’s test followed by Dunn’s test were utilised to test differences across different environmental compartments and types of sampling sites (reference vs tributary vs main river). Unique and shared OTUs were calculated to focus on the fine-scale variation within and between microbial communities.

For beta diversity, robust Aitchison distance was employed, as recommended by Martino et al. (2019), and graphically represented in a principal-coordinates analysis (PCoA) using the Euclidean dissimilarity index. Pairwise Permutational Multivariate Analysis of Variance (PERMANOVA) implemented in the R package pairwiseAdonis (Martinez-Arbizu, 2020) was used to assess dissimilarities among the three environmental compartments (water, biofilm, and sediment) and sampling site.

Putative ecological function profiles were generated applying Functional Annotation of PROkaryotic TAXa (FAPROTAX) implemented in microeco (Liu et al., 2021). FAPROTAX is a database that maps prokaryotic clades (e.g., genera or species) to established metabolic or other ecologically relevant functions, using the current literature on cultured strains (Louca et al., 2016). Putative ecological functions were determined to each environmental compartment and functions for abundances higher than 1%. We focused only in those ecologically relevant functions linked to the ecosystem service “degradation of chemicals”. Human and parasite-related functions were not considered in our analysis.

To explore the connections between beta diversity and environmental variables distance-based redundancy analysis (db-RDA) using Euclidian distance was employed (Legendre and Gallagher, 2001). The environmental variable matrix consisted of physicochemical parameters, nutrients, and antibiotic stress, represented as TU_MIC_. Prior to analysis, OTU tables and environmental variables were transformed using the robust Aitchison distance and Hellinger transformations, respectively, following Legendre and Gallagher (2001). The Variance Inflation Factors (VIF) for the constraining parameters were evaluated to determine if the constraints were redundant, and explanatory variables with VIF scores greater than 10 were excluded in order to build the “best” db-RDA model. The global significance of the “best” db-RDA model was evaluated via an analysis of variance (ANOVA; 999 permutations). All db-RDAs and subsequent permutation tests were conducted using the R package {vegan} (Oksanen et al., 2020).

DESeq2 (Love et al., 2014) was employed to perform differential abundance analysis (DAA). The negative binomial distribution was used to model the observed abundances, with data normalization performed using scaling methods to account for variations in sampling sites. The Wald test was used, with a parametric fit type, and the resulting p-values were subjected to multiple testing correction using the Benjamini and Hochberg method. OTUs demonstrating a fold-change of two or greater and adjusted p-values < 0.01 were considered statistically significant. All statistical analyses, data processing and visualisation were performed in R (R Core Team, 2021).

### 2.5. Normalisation of micropollutant concentrations

The measured environmental concentrations of 153 micropollutants, including antibiotics, detected at the sampling sites are reported in Inostroza et al., (2023b). This work describes in detail the analytical methodologies used, including sample collection, storage, extraction methods, LC- and GC-HRMS instrumentation, and the measured environmental concentrations of micropollutants in the River Aconcagua Basin. The data is accessible in Inostroza et al., (2023b) and through the open-access Zenodo repository (Inostroza et al., 2023b). We attempted to normalise the pollution profiles with available hazard information. However, we were only able to do so for antibiotics using minimum inhibitory concentration (MIC) data. For the rest of the micropollutants, we summed their measured environmental concentrations (CAChems) as surrogate metric. This latter approach overlooks the bioactivity of the micropollutants, but it provides a relative weight of the stress caused by micropollutants in the river basin.

The minimum inhibitory concentration (MIC) denotes the lowest concentration of an antimicrobial agent that inhibits the growth of a microorganism. MIC values were retrieved for each antibiotic from Bengtsson-Palme et al. (2016), who predicted size-adjusted lowest MIC also referred as “upper minimal selective concentration (MSC) boundaries”. The use of these MIC values allows to normalise measured antimicrobial environmental concentrations and weight the effect of antibiotics on bacterial communities. There are substantial evidence that concentrations below the MIC can lead to the selection of resistant bacteria (Andersson and Hughes, 2012; Gullberg et al., 2014, 2011; Liu et al., 2012). In fact, Gullberg et al. (2011) reported MSCs (i.e., the lowest concentration of an antimicrobial agent at which resistant strains have a competitive advantage) in the range of 1/230 to 1/4 of the MIC for different antibiotics.

To evaluate the impact of micropollutant including antibiotics on bacterial communities, we employed a component-based strategy. Specifically, we applied the dose or concentration addition (CA) mixture toxicity concept (Gustavsson et al., 2017; Inostroza et al., 2024; Spilsbury et al., 2020) using the following equation:

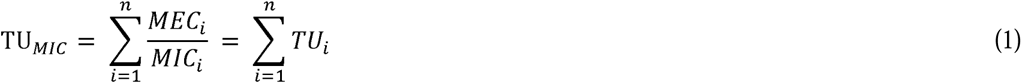

In equation (1), *MEC*_i_ represents the measured environmental concentration of antibiotic *i*, and *MIC*_i_ represents the corresponding minimum inhibitory concentration (MIC) of antibiotic *i*. The ratio *MEC*_i_/*MIC*_i_ provides a dimensionless measure of the individual toxicity contribution for each mixture component in a given sample, also known as a “Toxic Unit”.

All data curations including OTU tables, MIC concentrations, and data analysis scripts are openly available on Github (https://github.com/TO_BE_UPLOADED_ON_ACCEPTANCE) for further utilisation. Sequences obtained in this study were deposited at the National Center for Biotechnology Information (NCBI) under the project accession number PRJNA1080491.

## 3. Results and discussion

We assessed the diversity of bacterial communities sampled simultaneously from water, biofilms, and sediments in tributaries and the main river course of the River Aconcagua basin. Our study complements previous works that focussed on single compartments i.e., water (Li et al., 2023; Mansfeldt et al., 2020), biofilm (Aubertheau et al., 2017; Chonova et al., 2018; Freixa et al., 2020), sediment (Gao et al., 2019; Haller et al., 2011; Shao et al., 2023; Xie et al., 2016; Zhang et al., 2022), or on water and sediment together (Drury et al., 2013; Guo et al., 2016; Ibekwe et al., 2016; Valverde et al., 2021).

Sampling sites were selected based on differences in land use patterns and micropollution gradients (Inostroza et al., 2024), see also Figure S1. A maximum of 28 micropollutants were found at sites RS1, RS2, and RS3. These low numbers, accompanied by low concentrations, are a result of the low urbanisation in the area, supporting the use of these locations as “reference sites” (Inostroza et al., 2024). In contrast, up to 80 micropollutants were quantified at tributary sites and up to 71 in the main river course. The high number of micropollutants in tributary streams likely results from intensive agriculture and WWTP discharges near the sampling sites. Additionally, we determined significantly higher nutrient loads, conductivity, oxygen levels, antibiotic stress (TU_MIC_), and micropollutants (CAChems) in tributaries and the main river course (sites associated with agriculture and urban land uses) compared to reference sites (Figure S2).

We attempted to calculate toxic units for the encountered chemicals, i.e., to normalise their concentrations to their toxicity. Nevertheless, this turned out to be a major challenge, due to a critical lack of data on the bacterial toxicity of most of the chemicals found at the sampling sites. Appropriate quantitative structure-activity relationships (QSARs) are also not available for bacteria, so that *in silico* toxicity estimates were also not possible. Consequently, we were only able to assess the toxicity of antibiotics and chemicals with antimicrobial properties. This data was used to calculate antibiotic stress (TU_MIC_), analysed as a variable potentially influencing bacterial community structure.

### 3.1. Structure of bacterial communities from sites with different land uses

A total of 4,508,501 sequence reads were obtained and assigned to 16,018 operational taxonomic units (OTUs), comprising 7,878, 5,283, and 4,937 OTUs in sediment, biofilm, and water samples, respectively. The presence of singletons and doubletons accounted for only 290 (3.35%) and 125 (1.44%) sequences respectively, when all environmental compartments are considered. We assessed if removing or including singletons would affect alpha-diversity metrics i.e., Chao 1, which takes into account singletons and doubletons. The inclusion of low-frequency OTUs did not affect alpha- and beta-diversity analyses (Kruskal-Wallis’s test, *p*-adjusted < 0.001), therefore singletons and doubletons were not removed.

Distinct taxonomic patterns were observed at the phylum level across the different environmental matrices (i.e., water, biofilm, and sediment; Figure 1A). Proteobacteria exhibited a high median relative abundance across all matrices (30-55%), with significant higher abundances in the water compartment (Figure S3). Bacteroidetes had high median relative abundances primarily in water and biofilm (∼15% in each), whereas Actinobacteria showed significantly higher median relative abundance in sediment (25%) and Cyanobacteria in biofilms (40%) (Figure S3). The same phyla were also the most prevalent in sediment and water samples exposed to chronic pollution derived from fertiliser industry and mining (Valverde et al., 2021). In terms of the dynamic of these phyla across different sampling sites, Bacteroidetes had higher relative abundances along the main river course and tributaries which have been characterised by higher micropollutants concentrations (Inostroza et al., 2024, 2023b). Conversely, Cyanobacteria had lower abundances along the main river course and tributary sites in biofilm samples, which is consistent with previous studies (Aubertheau et al., 2017). Välitalo et al. (2017) concluded that Cyanobacteria are amongst the most sensitive microorganisms to antibiotics. We hypothesised that their abundances in the tributary and main river course are reduced (in comparison to samples from the reference sites) by the toxic pressure from antibiotics and antimicrobial chemicals emitted in WWTP effluents.

**Figure 1.**
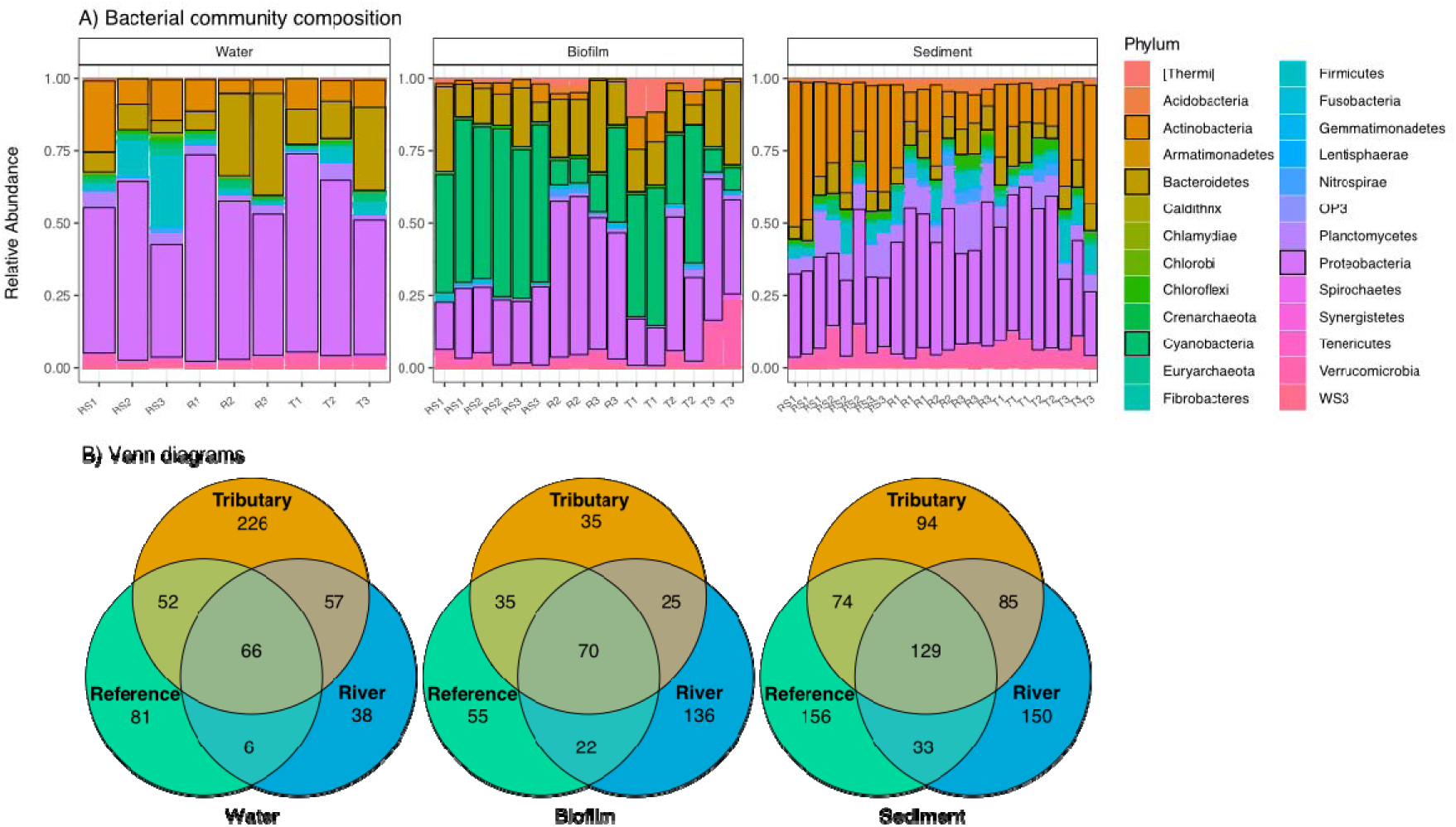
A) Relative abundance of bacterial phyla per sampling site and across environmental matrices: water (left), biofilm (centre), and sediment (right). Top phyla contributing significantly to at least one matrix is highlighted in black boxes in legend. B) Venn diagrams showing the number of unique and shared OTUs, measure of rare or microdiversity, between sampling sites per environmental compartment.

The assessment of unique and shared OTUs across samples revealed compartment-specific distributions. We detected 435 (5.0%), 345 (4.3%), and 226 (2.6%) unique OTUs in sediment, water, and biofilm respectively (Figure 1B). No clear pattern of consistently higher or lower diversity was observed across reference sites, tributaries, and samples from the main river (Figure 1B). Interestingly, the highest number of unique OTUs within the water compartment was found in tributaries, while the main river course exhibited the highest number of unique OTUs in biofilm communities. Sediment communities from reference sites demonstrated the greatest number of unique OTUs. This last finding aligns with previous studies that report higher numbers of unique OTUs in non-polluted sediments compared to their polluted counterparts (Haller et al., 2011).

### 3.2. Alpha diversity

Sediment-bacterial communities exhibited significantly higher alpha-diversities (described as Chao1- and Shannon-indices as well as eveness) than bacterial communities inhabiting water and biofilm, which had similar levels of alpha diversity (Dunn’s test, *p*-adjusted□<□0.05; Figure 2). These findings are consistent with previous studies, all showing that sediments harbour in general greater bacterial biodiversity (Lozupone and Knight, 2007; Valverde et al., 2021), and that water and biofilm habitats both display lower alpha diversity levels (Walters and Martiny, 2020). This pattern is driven by the greater heterogeneity of sediments, in comparison to water and biofilms, resulting in spatially/resource-driven niche partitioning, which increases bacterial alpha diversity (Gibbons and Gilbert, 2015).

**Figure 2.**
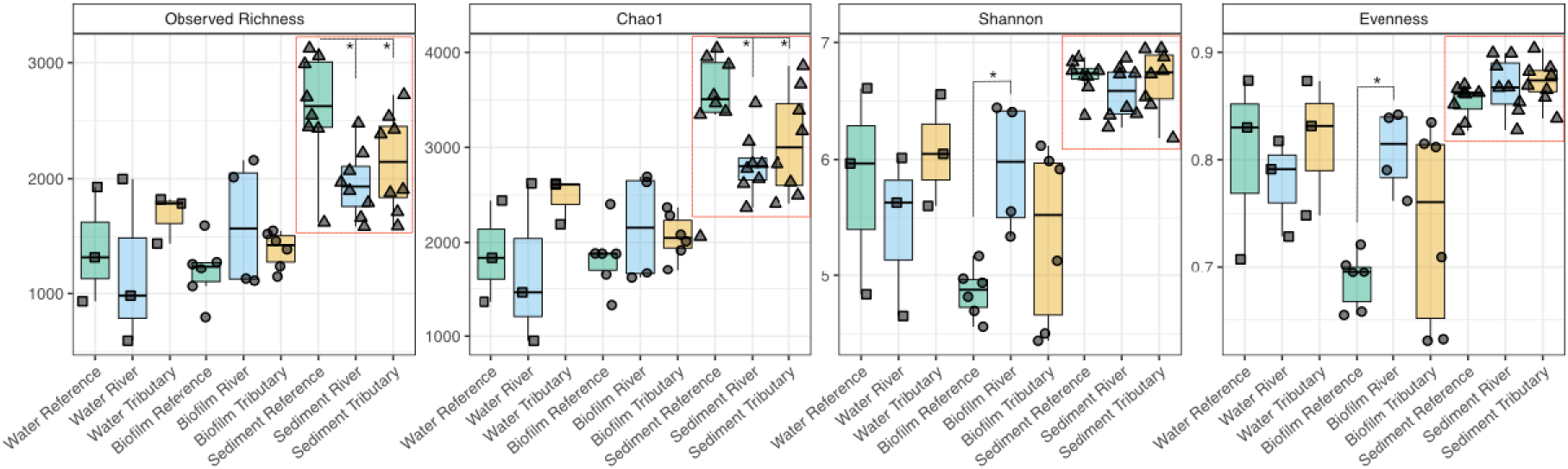
Alpha diversity metrics for bacterial communities in the River Aconcagua basin. Significant differences among environmental compartments and sampling sites are highlighted by a red box and asterisks, respectively (pairwise comparison using Dunn’s test, p-adjusted□<□0.05). Sampling sites are coloured coded (light green: reference sites, light blue river sites, light orange: tributary sites).

Sediment samples from reference sites harboured significantly higher alpha diversities compared to those under higher land use pressure, agricultural activities (tributaries) and WWTP effluent streams / urban areas (main river course), (Dunn’s test, *p*-adjusted□<□0.05). These latter sites were characterised by high micropollutant concentrations (Inostroza et al., 2023b). Conversely, biofilms retrieved from reference sites had the lowest median alpha diversity (Figure 2). Regarding the water compartment, the highest alpha diversity (median values) was observed in tributary sites (Figure 2), which were located in agricultural areas and small-sized urban areas with several discharges from WWTPs and high antibiotic stress (Figure S2). Our findings in the water compartment, specifically those sites affected by urban areas, are in line with previous studies (Ibekwe et al., 2016; Mansfeldt et al., 2020), which reported lower alpha diversity in surface water affected by WWTP discharges.

### 3.3. Beta-diversity and underlying factors influencing bacterial community structure

We identified significant dissimilarities between bacterial communities in water, biofilm, and sediment compartments (Figure S4; PERMANOVA test, *p*-adjusted < 0.001). Consistent with previous studies (Garner et al., 2016; Valverde et al., 2021), sediment bacterial communities displayed dissimilarities across different sampling sites with distinct pollution pressure (PERMANOVA test, *p*- adjusted < 0.001). Reference sites positively correlated with phosphate and light and negatively with nitrate, micropollutants (CAChems), and antibiotic stress (TU_MIC_) (Figure 3A). Tributary sites affected by agriculture (T2 & T3) correlated negatively with phosphate, silicate, light, pH, and temperature. However, tributary site T1, affected by agriculture and WWTP’s discharges, correlated with nitrate and TU_MIC_ (Figure 3A). Lastly, the diversity of bacterial communities sampled along the main river course, that is affected by rural (R1) and urban areas (R2), correlated with TU_MIC_ and the river site R3, close to the river debouch, negatively correlated with pH and temperature. (Figure 3A). Overall, antibiotic stress (TU_MIC_) was a major variable explaining the variations in the bacterial community for the river sites R1 and R2 and tributary site T1, consistent with the presence of antibiotics recently reported throughout the river basin (Inostroza et al., 2023a).

**Figure 3.**
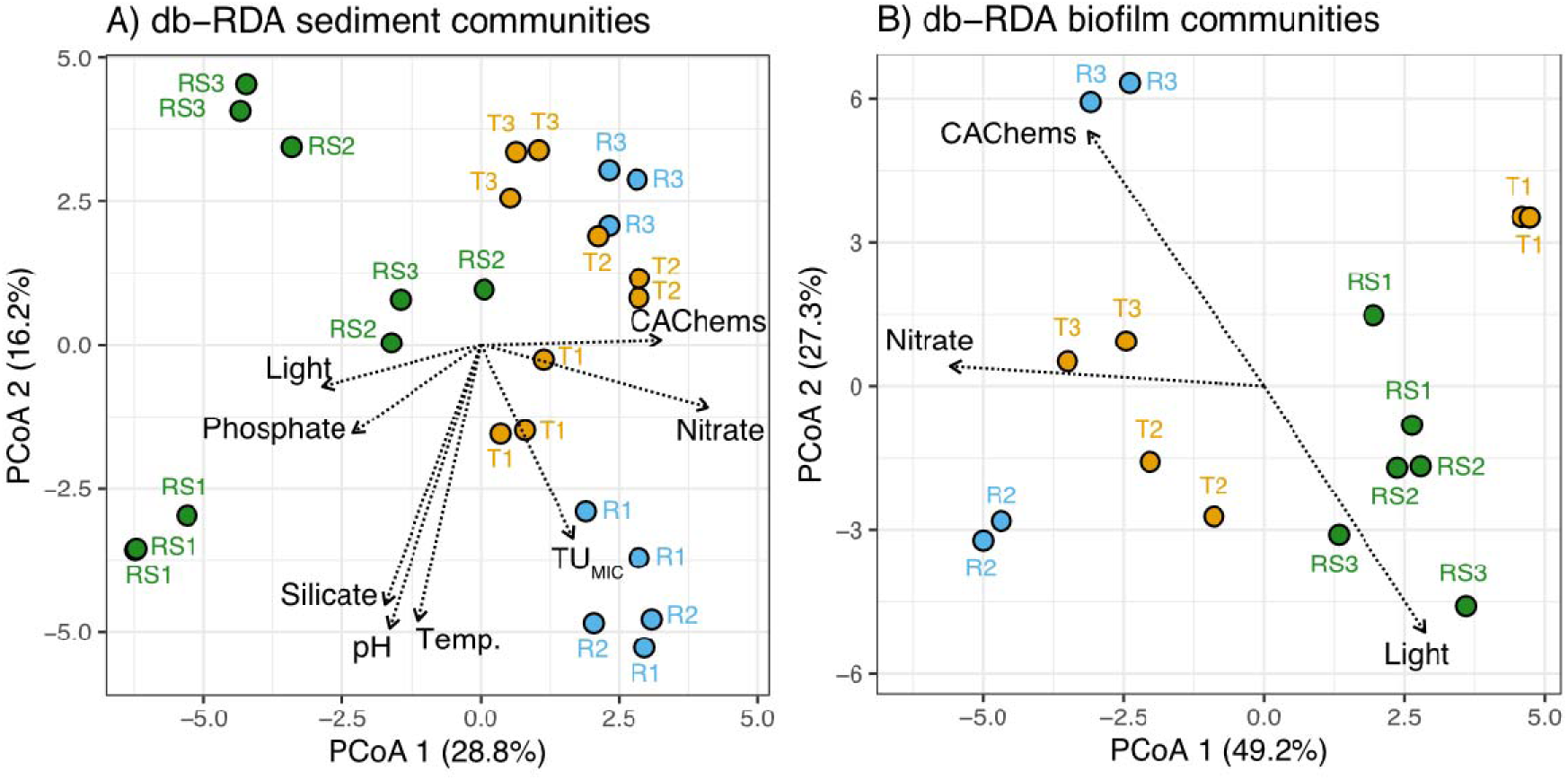
Distance-based redundancy analysis (db-RDA) showing the significant models (ANOVA like permutation test for db-RDA, *p*-value□<□0.001) containing environmental parameters, including nutrients (i.e., nitrate, phosphate, and silicic acid), physico-chemicals variables (i.e., light, temperature, and pH), antimicrobial stress (TU_MIC_), and summed measured concentration of micropollutants (CAChems). These variables significatively explained the observed dissimilarities in bacterial communities in A) biofilm and B) sediment samples.

We estimate the highest TU_MIC_ in sampling sites that are influenced by WWTP discharges (R2 & R3) and particularly in R1, a rural but still populated area lacking WWTP facilities. Indeed, R1 exhibited the highest concentrations of erythromycin and clarithromycin (8.7 ng/L and 10.5 ng/L (Inostroza et al., 2023a)), suggesting the discharge of untreated wastewater and/or the application of WWTP sludge to areas upstream of R1. This finding is consistent with earlier studies identifying high concentrations of antibiotics in rural areas without WWTPs (Gray et al., 2020; Yang et al., 2020). Furthermore, non-domestic effluents related to mining camps and animal farming located far upstream of R1 may represent an additional source of antibiotics.

Although we observed a positive correlation between TU_MIC_ and river and tributary sites (R1, R2, T1), TU_MIC_ did not correlate with micropollutant concentrations (CAChems). Thus, they cannot be considered a proxy for the complex mixture of pesticides, other pharmaceuticals, personal care products, and other organic synthetic chemicals co-occurring at low concentrations in the River Aconcagua basin. Because TU_MIC_ is comprised of only a few antibiotics, we hypothesise that the suggested metric may be representative of other antibiotics and/or antimicrobial chemicals overlooked by the target list used in Inostroza et al., (2023a).

The impact of antibiotic stress was evaluated as abundance changes at the genus level in bacteria inhabiting sediment (Figure 4), which was the only compartment significantly affected by antibiotics (Figure 3). The full list of genera, along with their respective fold-change direction, and references, is provided in Table S3. Communities sampled at river site R1 showed the highest increase in the number of genera compared to reference sites (Figure 4A), followed by R2 (Figure 4B). River site 3 (R3) was not correlated with antibiotic stress (Figure 3A).

**Figure 4.**
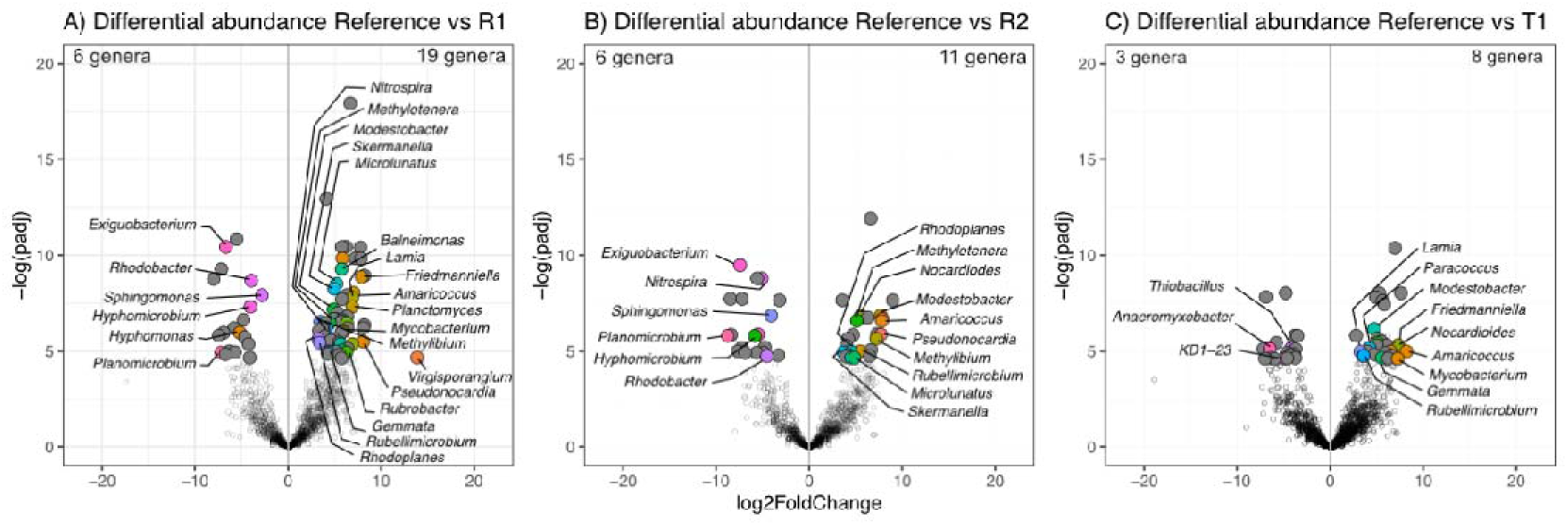
Differential abundance analysis of sediment communities impacted by antimicrobial stress (TU_MIC_). A) Comparison between the impacted site R1 and the nearest reference site RS1. B) Comparison between the impacted site R2 and the nearest reference site RS2, and C) between tributary T1 and RS1. Log_2_foldChange represents the change in abundance (reduction or increase). Only genera showing significant changes are provided in colour, and total numbers are presented. Each colour represents one specific genus, which are provided in Table S3.

In total, up to 19 genera responded to antibiotic stress, either increasing or decreasing their abundances (Figure 4). Most of these genera have been characterised as bacteria with the capacity to biodegrade organic chemicals in polluted sediments, such as ether-containing pollutants (Simon et al., 2006), pesticides (Garg et al., 2021), polycyclic aromatic hydrocarbons (Uhlik et al., 2012), and other micropollutants (Brunhoferova et al., 2022). However, not only bacteria known to be involved in the degradation of toxic chemicals were affected by antibiotic stress. *Nistrospira*, known to encode multidrug resistance (Liu et al., 2019), showed a significant increase in sediments from R1. On the other hand, antibiotic-resistant bacteria such as *Exiguobacterium* and *Planomicrobium* (Gao et al., 2012), were adversely affected by antibiotic stress in R1 and R2. In biofilms, we also observed increased abundances of bacteria known to degrade micropollutants (Brunhoferova et al., 2022; Xu et al., 2008), other organic pollutants including alkanes, pyridine, phenols, and phenanthrene (Park et al., 2020; You et al., 2023). Even though the levels found in the environment are in general below the minimum inhibitory concentration for most bacteria, they may still result in an increased selection pressure at some sites (Kristiansson et al., 2011). Furthermore, it has been shown that antibiotic concentrations below minimum inhibitory concentrations (MICs) can select for resistant bacteria (Berglund et al., 2014; Gullberg et al., 2011).

Bacterial communities in biofilm samples collected at tributaries and main river sites were not significantly different, although both were significantly different from reference conditions (PERMANOVA test, *p*-adjusted < 0.05). The observed dissimilarities were better explained by different light conditions in reference sites, nutrient, particularly nitrate, in tributary site T2, and micropollutant loads (CAChems) in the main river site R3 (Figure 3B). Lastly, we did not observed dissimilarities in the water compartment (PERMANOVA test, *p*-adjusted > 0.1), in line with previous studies (Valverde et al., 2021), likely due to the high degree of connectivity of this compartment.

### 3.4. Potential ecological changes in bacterial communities

We assessed potential functional shifts in bacterial communities across different sampling site types (reference, tributary, and main river sites) and environmental compartments (water, biofilm, and sediment). We used FAPROTAX, which contains an extensive list of ecologically relevant bacterial functions (Louca et al., 2016). Nevertheless, we focused on functions related to the degradation of organic chemicals and plastic, as these serve as proxies for the ecosystem service of degrading toxic chemicals in aquatic environments.

Our findings indicate mixed bacterial degradation responses dependent on the environmental compartment. In water and biofilm, we observed a reduction of hydrocarbon degradation function in tributaries and the main river course compared to reference sites. However, tributaries in the sediment compartment showed a significant increase in this function (Dunn’s test, p-adjusted < 0.05, Figure 5). Similarly, the aromatic compound degradation function decreased in tributaries and main river course sites in water and sediment compartments compared to reference conditions (Dunn’s test, p-adjusted < 0.05, Figure 5). Similar to our findings, Yao et al. (2022) reported mixed responses in bacterial communities of polluted sediments. They observed a decline in aromatic compound degradation and an increase in hydrocarbon degradation compared to reference conditions. Conversely, no clear pattern emerged in biofilm samples. Our study suggests a potential promotion of plastic, aromatic and aliphatic hydrocarbons in biofilm communities at tributary sites (Figure 5).

**Figure 5.**
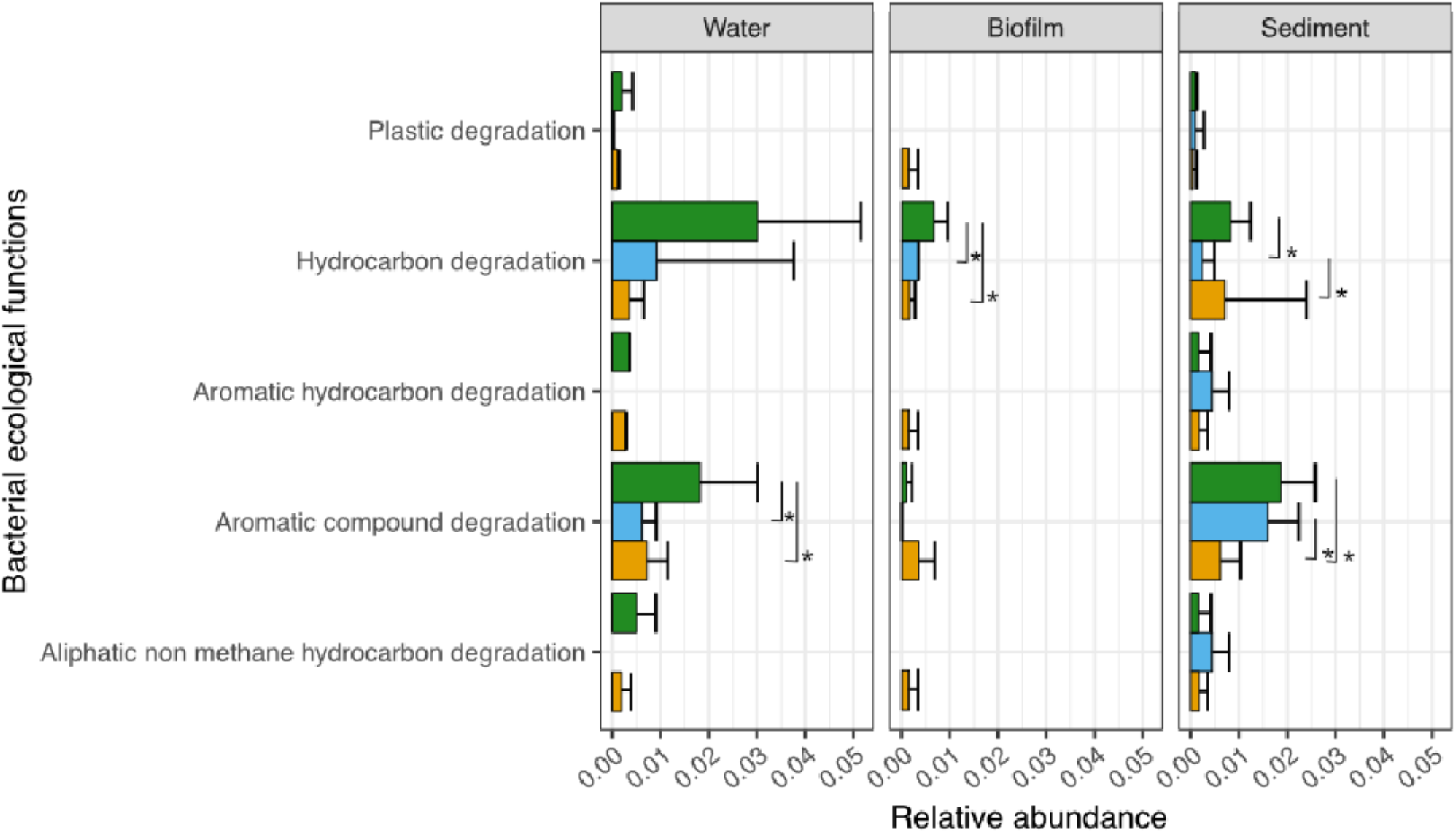
Functional bacterial community profiles of each environmental compartment in different land uses based on FAPROTAX. Significant differences (Dunn’s test, p < 0.05) are marked with an asterix. Sampling sites are coloured coded (green: reference sites, light blue river sites, orange: tributary sites).

The observed functional reductions might be attributed to pollution pressure from the micropollutant loads detected in tributaries and the main river course of the River Aconcagua basin (Inostroza et al., 2023b). Furthermore, a recent mixture risk assessment study predicted that tributaries and the lower part of the main river course are at risk due to pollution pressure from mixtures of pesticides, pharmaceuticals, and industrial chemicals in the same basin (Inostroza et al., 2024). It is important to note, this mixture risk assessment was based on ecotoxicological information from the most sensitive trophic levels (i.e., algae, crustaceans, and fish) and does not incorporate impacts on bacterial biodiversity.

## 4. Conclusions

Bacterial communities inhabiting water, biofilm, and sediments displayed significantly distinct structures. Sediment communities exhibited the highest alpha-diversity, especially in reference sites, aligning with their greater environmental and resource heterogeneity. In water and biofilm, we observed mixed responses: biofilm communities increased their alpha-diversity in polluted sites (e.g., nutrients and organic chemicals), while bacteria in the water compartment showed their highest alpha- diversity in tributaries characterised by high nutrient loads.

Beta-diversity analyses underscored the influence of distinct site-specific environmental stressors, including nutrients, micropollutants, and antibiotics for some river sediments and nutrients and micropollutant loads in biofilms. Notably, some sites exhibited increased levels of bacterial genera known for pollutant degradation, highlighting the potential for functional adaptability under pollution pressure.

Our findings reveal a negative impact on ecosystem services associated with the bacterial degradation of organic chemicals in regions of the River Aconcagua basin affected by high pollution loads (tributaries affected by agriculture and the main river course by mixtures of pesticides and pharmaceuticals from WWTP discharges). However, sediment samples from tributary sites unexpectedly exhibited increased hydrocarbon degradation potential. This finding prompts further investigation into the dual role of sediments as both sinks and sources of pollutants and their transformation products. Additionally, we suggest experimental functional approaches to determine whether such predictions accurately reflect reality through controlled experiments.

Our study is based on a single sampling campaign, a clear shortcoming, limiting our findings to a unique season. We recognise the necessity of capturing seasonal dynamics of microbial communities in relation to nutrient and antibiotic loads though additional sampling efforts.

Furthermore, our study underlines the critical need for empirical toxicity data on microbial organisms to safeguard essential ecosystem functions that bacteria provide. These would also allow to develop meaningful QSAR models for this organism group It remains crucial to incorporate bacterial ecotoxicological data for conducting comprehensive environmental risk assessments. However, we are concerned that similar challenges may arise when addressing other microorganism groups, such as fungi and/or micro-eukaryotes, which are also largely overlooked in the context of environmental risk assessment.

## 5. CRediT authorship contribution statement

**Pedro A. Inostroza**: Conceptualization, Methodology, Software, Validation, Formal analysis, Investigation, Data curation, Writing – Original Draft, Writing – review & editing, Visualisation. **Gerdhard L. Jessen**: Methodology, Writing – review & editing. **Feilong Li**: Methodology, formal analysis, Writing – review & editing. **Xiaowei Zhang**: Resources, Writing – review & editing. **Werner Brack**: Resources, Writing – review & editing. **Thomas Backhaus**: Conceptualization, Formal analysis, Data, Writing – Original Draft, Writing – review & editing, Funding acquisition.

## 6. Acknowledgments

We thank Alba Lopez Mangas and Monica del Aguila for fieldwork support. This work was supported by the FRAM Centre for Future Chemical Risk Assessment and Management at the University of Gothenburg, Sweden. Additional support came from The National Fund for Scientific and Technological Development of Chile (FONDECYT) Grant 1200252 (The National Research and Development Agency of Chile, ANID Chile) and COPAS COASTAL ANID FB210021 to GLJ.

## Supplementary Information

### SI Figures

**Figure S1.**
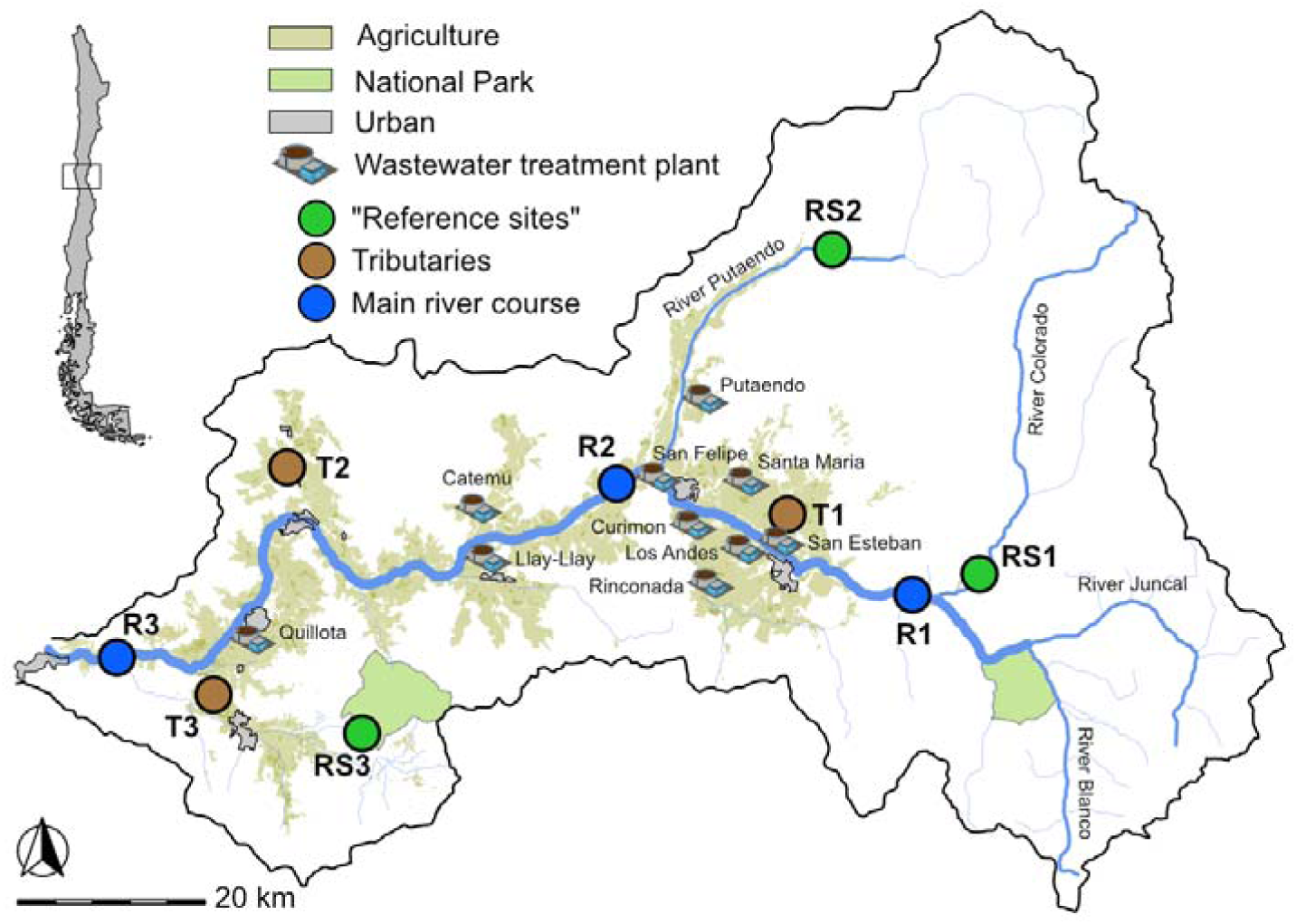
Location of sampling sites through the River Aconcagua Basin. Sites featuring “low” urban/agriculture pressures (“Reference sites”) in green, sites running through agricultural areas in brown, and sites from the main course of the river in light blue. Selected land use (agriculture, urban areas, and national parks) and location of WWTPs are showed in the figure.

**Figure S2.**
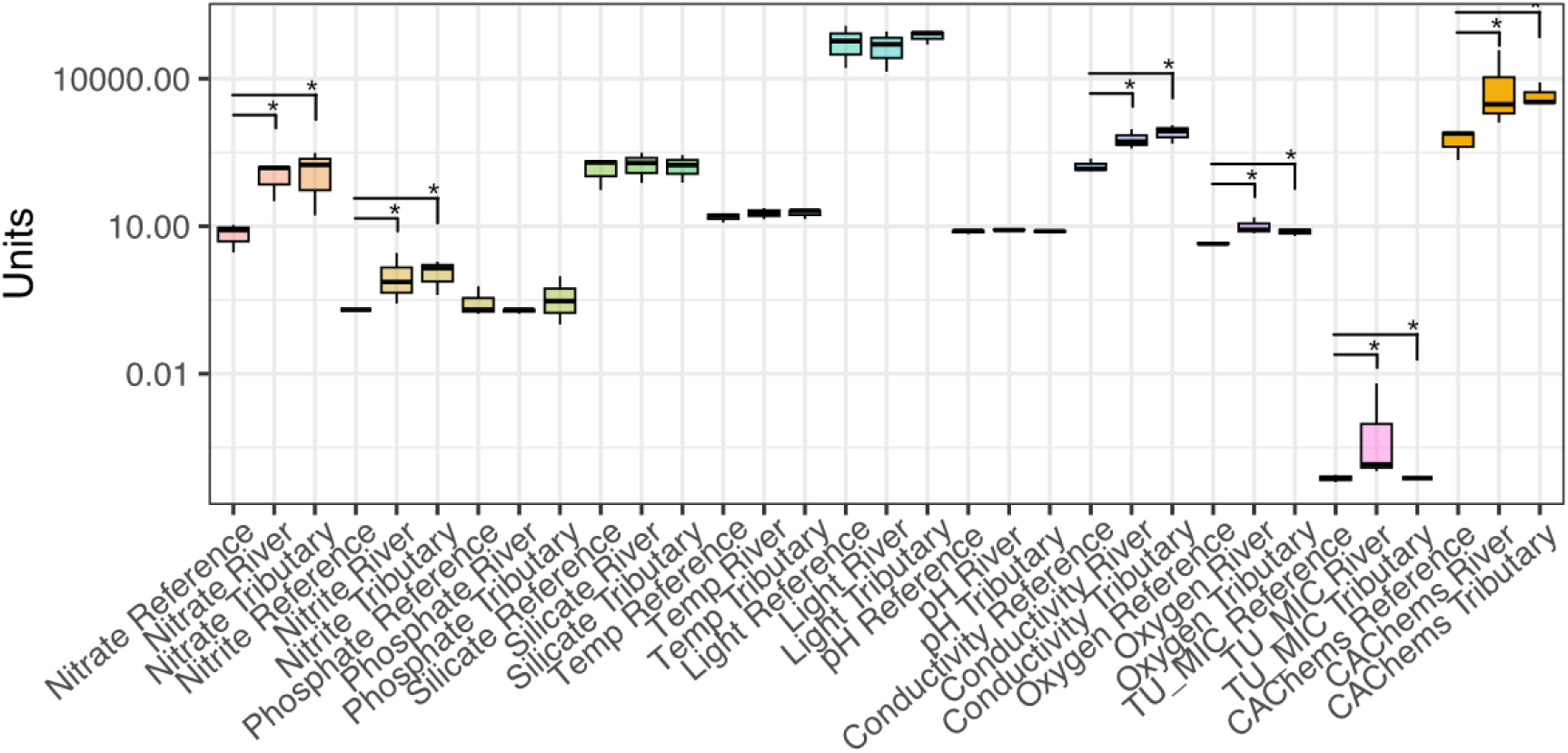
Environmental parameters in the River Aconcagua basin. Significant differences between reference sites and sampling site types are represented with the symbol *.

**Figure S3.**
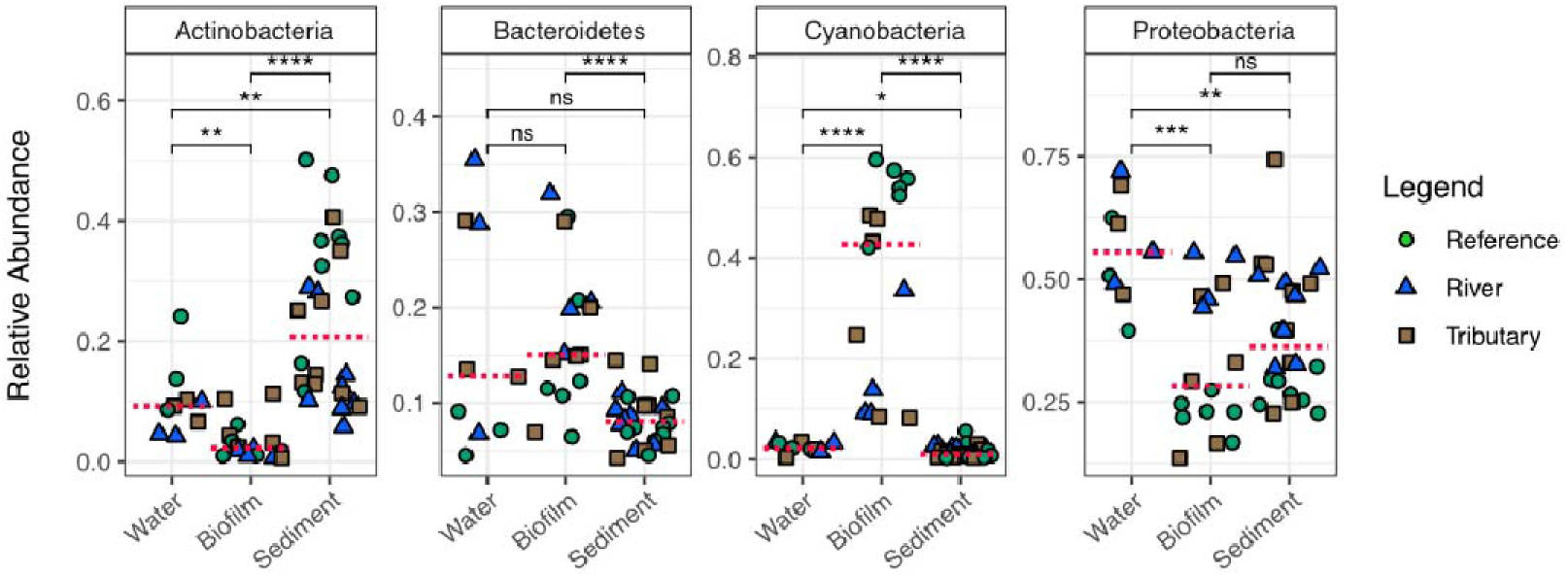
Bacterial composition of TOP-4 phyla in the River Aconcagua basin. Dashed red crossline represent the median in each of the environmental compartment. Significant level: 0.05 (*), <0.01 (**), <0.001 (***), <0.0001 (****), and n.s. = non-significant.

**Figure S4.**
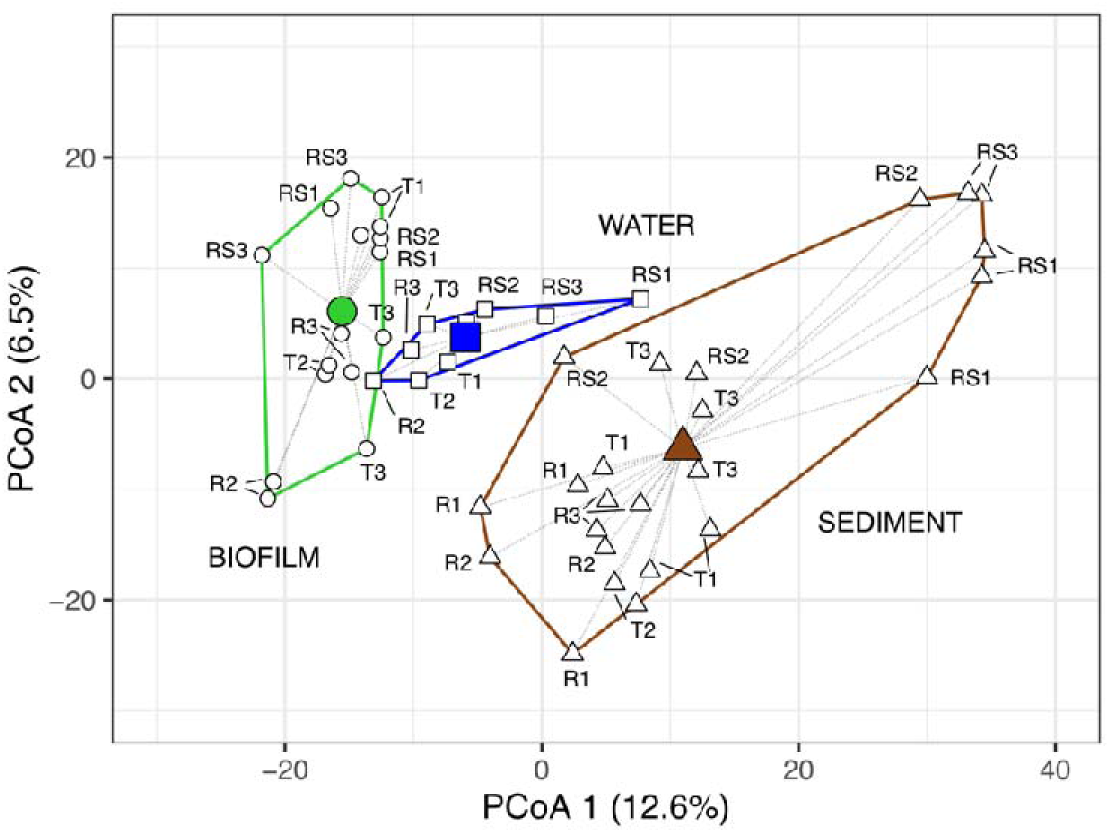
Degree of dissimilarity (beta diversity) between the studied environmental compartment at the River Aconcagua basin. The three compartments are significantly different from each other (PERMANOVA test, p-adjusted < 0.001).

### SI Tables

**Table S1.**
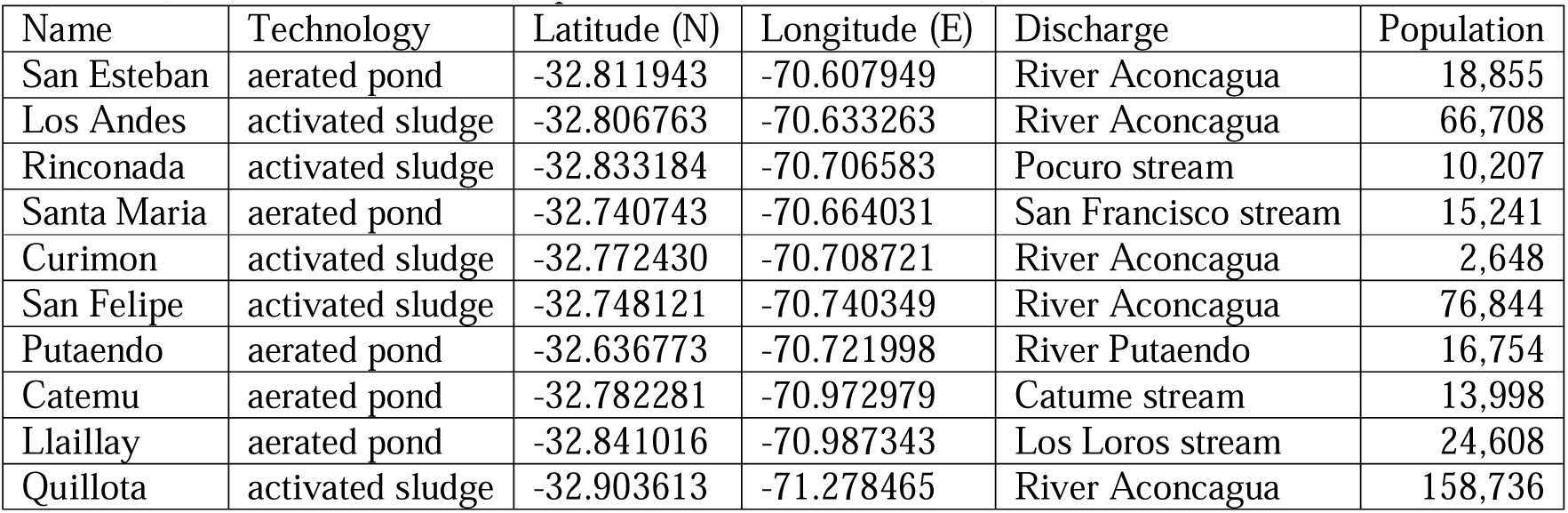
Information of WWTPs across the River Aconcagua basin. Geographic coordinates in decimal degree (WGS84). Updated serving population retrieved from the National Statistics Institute of Chile (INE) online database (http://resultados.censo2017.cl/).

**Table S2.** Sampling site information. Geographic coordinates in decimal degree (WGS84). WWTP= wastewater treatment plant.

| Site ID | Type | Latitude (N) | Longitude (E) | Odour | Colour | WWTP discharges | Floating material | Land use-type |
| --- | --- | --- | --- | --- | --- | --- | --- | --- |
| RS1 | “reference” | -32.854358 | -70.390044 | none | colourless | No | none | Reference |
| RS2 | “reference” | -32.509769 | -70.452537 | none | colourless | No | none | Reference |
| RS3 | “reference” | -33.003752 | -71.126355 | none | colourless | No | none | Reference |
| T1 | Tributary | -32.765909 | -70.613844 | farm | gray/brown | No | organic material | Agriculture |
| T2 | Tributary | -32.695661 | -71.212179 | sewage | colourless | Yes | debris | Agriculture |
| T3 | Tributary | -32.938952 | -71.329491 | sewage | colourless | Yes | organic material, bloom | Agriculture |
| R1 | Main river | -32.852416 | -70.502894 | none | gray | No | foam | Rural |
| R2 | Main river | -32.762305 | -70.839624 | none | colourless | Yes | foam | Urban |
| R3 | Main river | -32.916946 | -71.425322 | none | colourless | Yes | organic material | Urban |

**Table S3.**
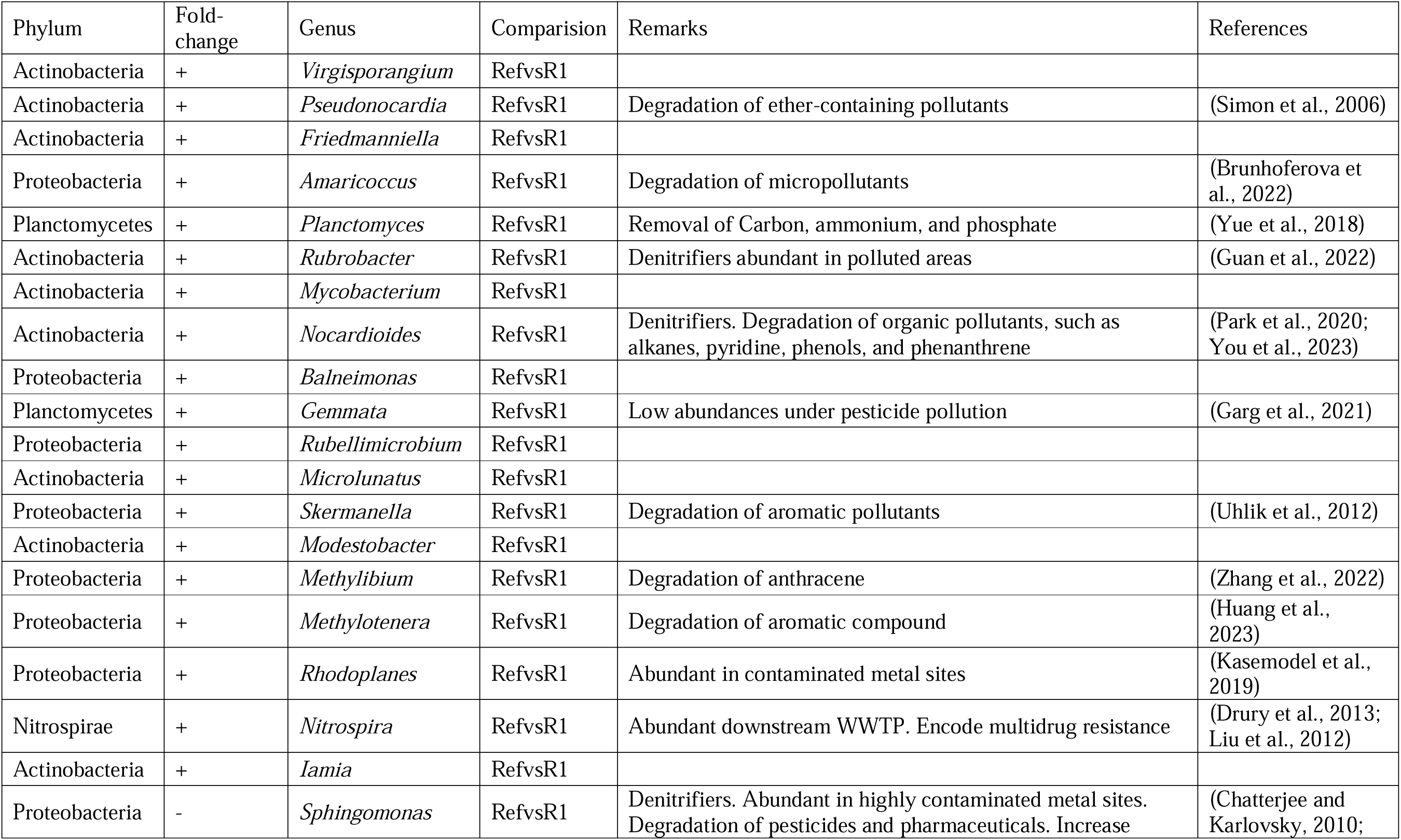
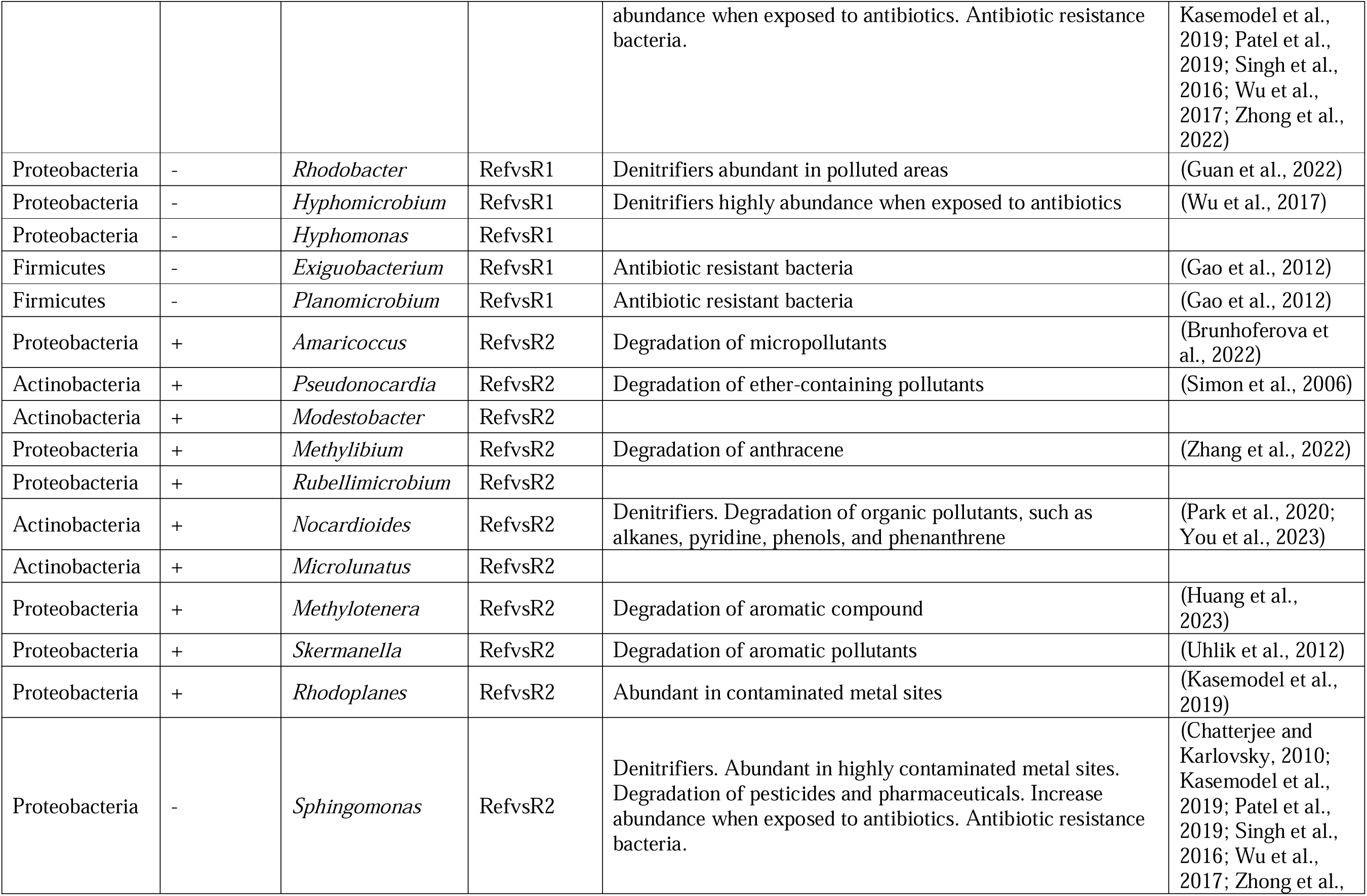
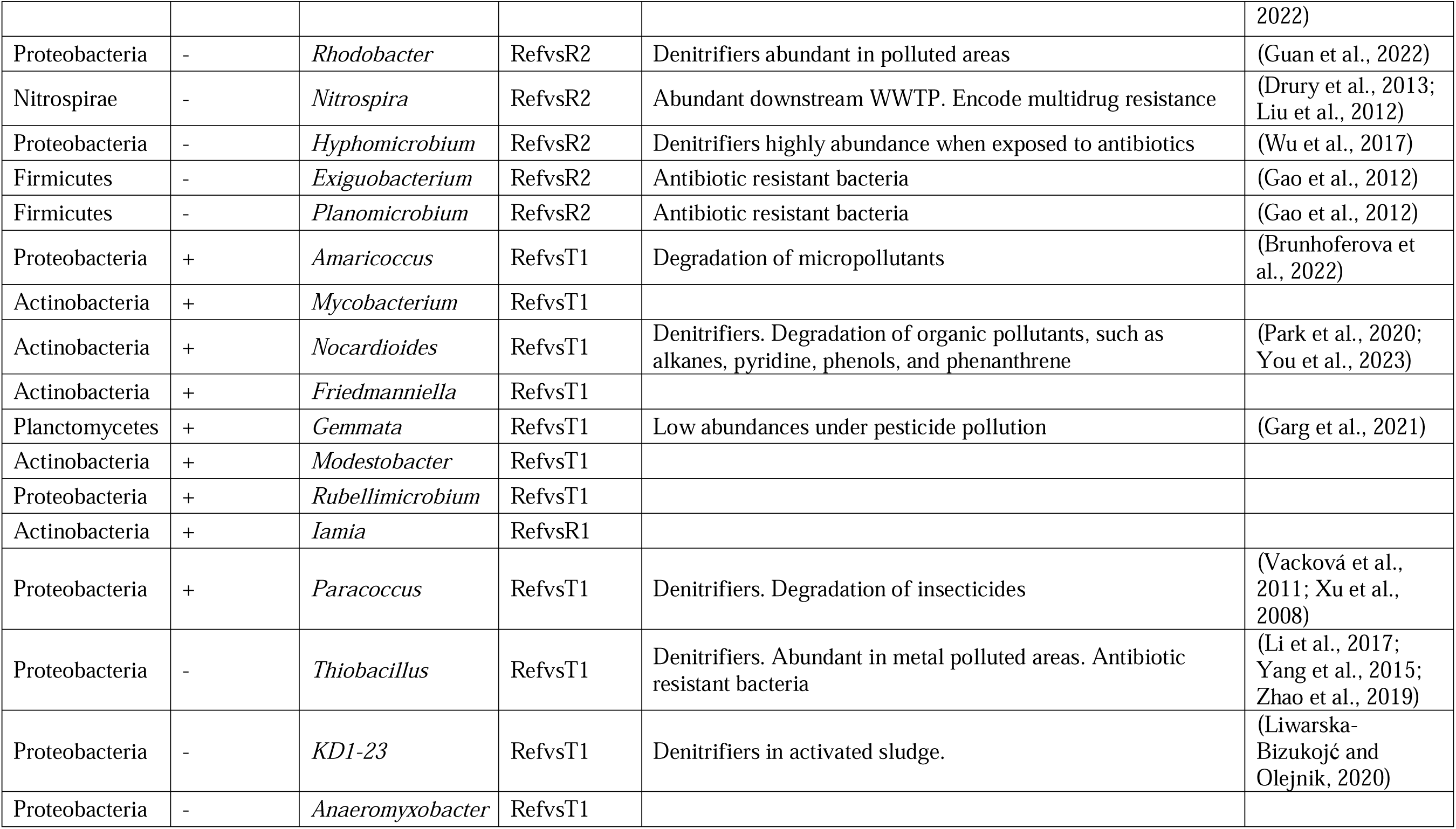
Differential abundance analysis summary table including phyla, direction of the fold-change, genera, comparision, ecological functional remarks, and references.

## Notes

### Competing Interest Statement

The authors have declared no competing interest.

